# ZipStrain Enables Rapid and Precise Strain-Resolved Metagenomics

**DOI:** 10.64898/2026.05.20.726564

**Authors:** Parsa Ghadermazi, Joanne B. Emerson, Matthew R. Olm

## Abstract

Strain-resolved metagenomics characterizes microbial communities at nucleotide-level resolution, enabling researchers to differentiate identical from closely related organisms and characterize population structure and gene content variation. Here we introduce ZipStrain, a program that performs highly accurate strain-resolved metagenomics over 500× faster than available methods while offering superior RAM management. Applied to a dataset of 2,754 samples spanning human populations, we identify a strain-sharing gradient across social relationships, reveal striking variation in clonal structure across bacteria and bacteriophage, and pinpoint genes whose nucleotide identity deviates from genome-wide expectations. ZipStrain is distributed as an open-source Python package and accompanying Nextflow pipeline at https://github.com/OlmLab/ZipStrain.

## Background

Shotgun metagenomics involves sequencing DNA from entire microbial communities, enabling simultaneous characterization of thousands of taxa within a single sample. Each taxon is itself a population of millions of cells that can differ by as little as a single nucleotide, and each of these cells is sequenced concurrently. Most currently used analysis tools handle this complexity by smoothing over fine-scale diversity to classify community composition at the species level (1,2). Methods capable of resolving this within-species diversity, often referred to as “strain-resolved” approaches, have yielded substantial insights inaccessible to species-level analyses. In the human gut such methods have illuminated patterns of mother-to-infant strain transmission (3), household strain sharing (4), and strain exchange in daycare settings (5). Beyond the human gut, strain-resolved metagenomics has advanced understanding of microbiome assembly and stability (6,7) and engineered microbiomes (8,9), microbial ecology and evolution (10,11) and clinical applications like monitoring strain engraftment following microbiome-based interventions (12).

Among the possible applications of strain-resolved metagenomics, determining whether a strain transmission event has occurred between two samples (e.g. from a mother to her infant) stands out as one of the most biologically and clinically consequential. Detecting strain transmission involves identifying highly similar genomes across multiple samples. This is complicated by the fact that closely related strains are frequently found in unrelated individuals, not because of direct transmission, but because of the global spread of human-adapted lineages (13,14). Two individuals may therefore harbor nearly identical strains acquired entirely independently, and distinguishing true transmission from such coincidental similarity requires extremely stringent genome-wide identity thresholds (e.g., 99.999% average nucleotide identity (ANI)). While these thresholds can accurately differentiate transmission from coincidental similarity (15), achieving the precision needed to apply them is difficult given the complex within-population diversity inherent to metagenomic samples.

Existing strain-resolved metagenomic approaches differ substantially in how they balance computational efficiency, resolution, and scalability. K-mer–based methods, such as Strainer and LongTrack (16,17), assign each genome a set of unique k-mers and achieve substantial computational speed. However, these approaches rely on obtaining high-quality genomes for strains across all samples to define unique k-mers, which is both difficult to achieve in complex environments and makes accuracy both sensitive to dataset composition and variable across studies. Marker gene–based methods, including StrainPhlAn (7), map metagenomic reads to a subset of conserved genes and calculate the similarity of these regions across samples. Although widely used, these methods consider only a fraction of each genome (typically around 1-10% of the total genome length) which may overlook biologically meaningful variation outside the marker gene loci. Further, these methods typically do not account for within-sample population diversity (15). Tools like InStrain (15) and MIDAS (6) adopt a genome-wide strategy, enabling detailed nucleotide-level comparisons across samples. However, as the number of samples or reference genomes increases, exhaustive pairwise comparison rapidly becomes computationally prohibitive. Further, most programs do not extend substantially beyond SNP calling and ANI calculation, leaving much of the information in genome-wide nucleotide-resolution data underutilized.

Here we present ZipStrain, a strain-resolved metagenomics tool that is faster and more flexible than existing methods without compromising accuracy. ZipStrain follows the general workflow of inStrain (15), but is built from the ground up using high-performance libraries such as Polars, DuckDB (18,19), and PyTorch (20) and exposes richer intermediate data for custom analyses. These design choices yield orders-of-magnitude gains in speed and efficiency, enabling comparisons across thousands of samples and large genome databases that were previously impractical or impossible. ZipStrain also introduces gene-level comparisons and identity-by-state (IBS) analysis (21). By applying ZipStrain to a global cohort, we show that strain-sharing tracks social connectivity, clonal structure varies across taxa, and that genes with evidence of *in-situ* selection can be identified. Together, these results demonstrate how removing the trade-off between resolution and scale enables new insights into microbial transmission and population structure.

## Results

### ZipStrain Workflow and Performance

ZipStrain utilizes a reference genome database and metagenomic mapping-based workflow **(Figure 1)**. Reads are first mapped to a reference genome database that can either be provided by the user or automatically constructed by ZipStrain (**Figure 1A**) using Sylph (22) and GTDB (23–27) (see methods for details). The resulting alignments are processed into per-position nucleotide frequency tables with integrated gene annotations and stored in an optimized, compressed format.

**Figure 1.**
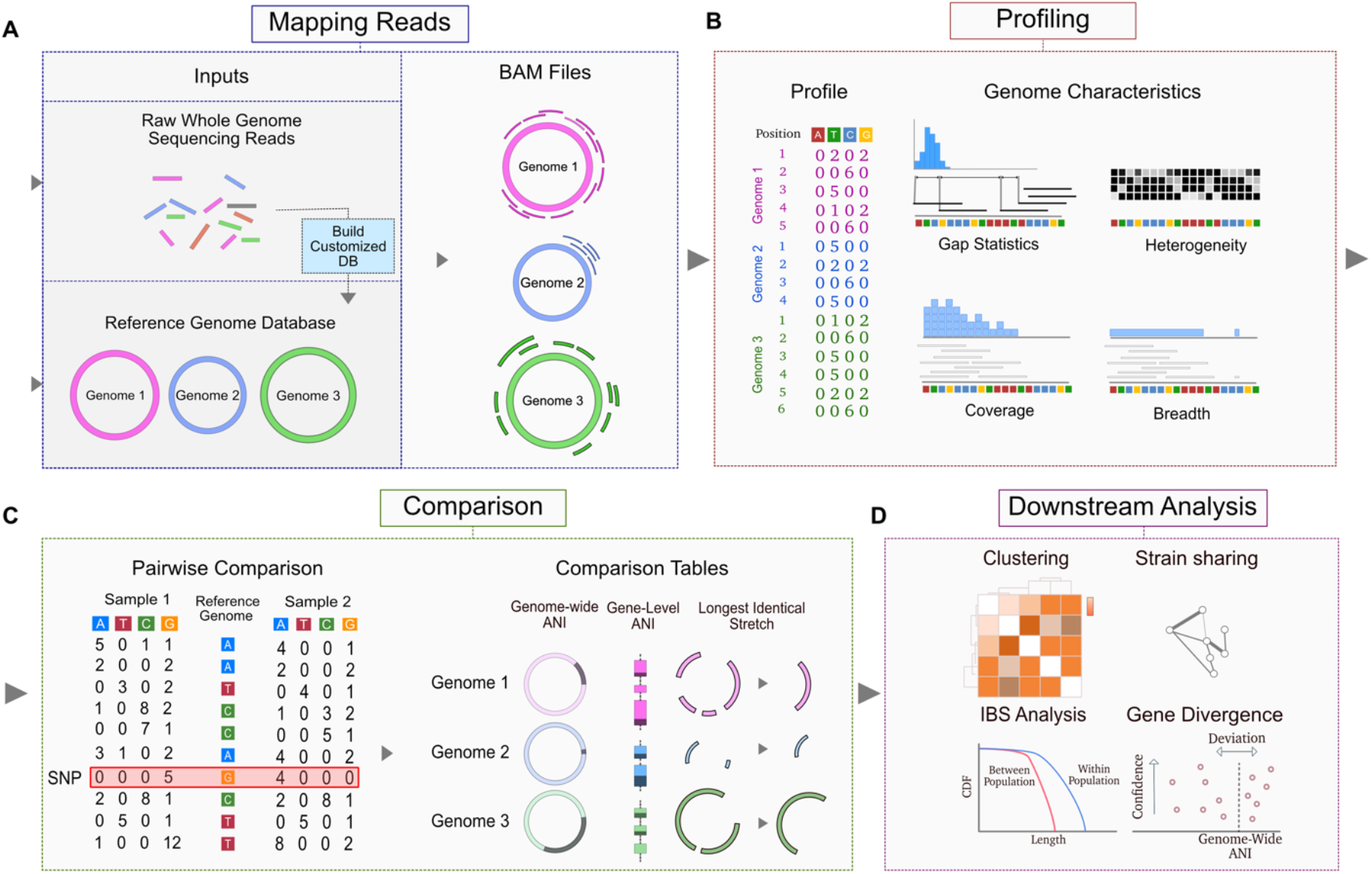
Overview of the ZipStrain workflow. Raw metagenomic reads are mapped to a reference genome database using a short-read aligner (e.g., Bowtie2), generating BAM files that are subsequently processed with the ZipStrain profile workflow. The reference genome database is either provided by the user or built by ZipStrain using tools that infer taxonomic composition of metagenomics samples from raw metagenomics reads. Profiling produces two outputs: **(A)** a nucleotide-resolution profile containing base frequencies along with gene and genome annotations at all covered positions, and **(B)** a genome characteristics file summarizing genome-level metrics such as coverage, breadth, number of mapped reads, read gap statistics, and strain heterogeneity. **(C)** Profiles from multiple samples can then be compared at either the gene or genome level. Genome-level comparisons generate metrics including genome-wide average nucleotide identity (ANI), number of identical genes, and the longest identical stretch between samples for each genome. **(D)** These comparison tables form the basis for downstream analyses in ZipStrain, including clustering, strain sharing, identity-by-state analysis, and gene-level divergence assessments.

Genome-level statistics are then calculated and presented for each genome in each sample, enabling accurate abundance estimation and presence/absence inference (discussed in detail below) (**Figure 1B**). The genome comparison workflow in ZipStrain compares all genomes across all sample pairs, producing a comprehensive pairwise comparison database (**Figure 1C**). Reported data from pairwise comparisons includes genome-wide average nucleotide identity (ANI), the number of positions at or above the minimum coverage threshold, the genes present in both samples, the number of genes with 100% ANI between the two samples, and the longest contiguous stretch of perfect sequence identity between the two samples for each genome. The latter two metrics are especially useful for evolutionary analyses, where the fraction of identical genes provides a proxy for the recency of clonal common ancestry and the longest identical tract reflects the age of the most recent horizontal gene transfer event between two strains (discussed in detail below) (21). By default, ANI is calculated using the popANI metric (15), though ZipStrain also supports conANI and a novel cosANI metric (see documentation for details). Together the ZipStrain outputs support a wide range of downstream analyses, several of which are described in the sections that follow (**Figure 1D**).

#### The Computational Cost of ZipStrain

ZipStrain achieves profiling and comparison speeds that are orders of magnitude faster than InStrain, while also significantly improving memory management (**Figure 2A–C**). To benchmark both tools, we profiled and compared 100 samples across single-genome and all-genomes modes. In the profiling workflow, ZipStrain was **16 ± 4×** faster than InStrain while also consuming less RAM (**Figure 2A**).

**Figure 2.**
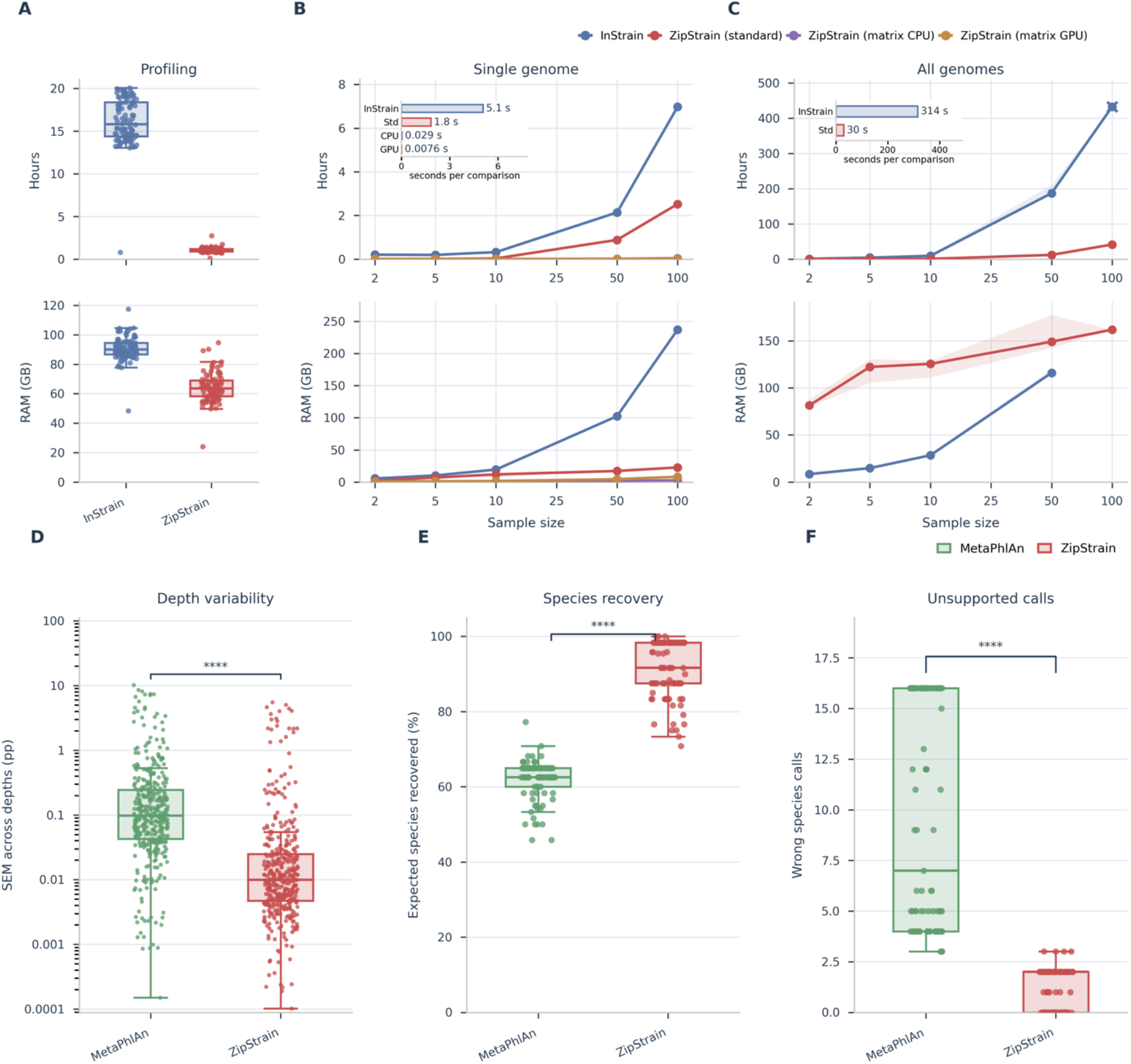
ZipStrain improves profiling performance and taxonomic consistency relative to existing tools. **A**, Runtime and peak RAM for profiling 100 metagenomic samples with InStrain and ZipStrain. Points show individual samples; boxes show the median and interquartile range. ZipStrain reduced profiling runtime by **15.95 ± 4.30-fold** relative to InStrain across paired samples (**mean ± s.d.; n = 100**). **B, C**, Runtime and peak RAM scaling for pairwise comparisons across sample sizes for **single-genome** (**B**) and **all-genome** (**C**) comparisons. Lines show median runtime or RAM and shaded bands show the interquartile range across replicates. Insets show runtime per comparison at **n = 100** in seconds. At n = 100, ZipStrain standard mode was **2.77-fold** faster than InStrain for single-genome comparisons, while matrix CPU and GPU modes were **175-fold** and **664-fold** faster, respectively. For all-genome comparisons, ZipStrain standard mode was **10.44-fold** faster than InStrain. **D**, Taxonomic abundance stability across sequencing depths, measured as the standard error of the mean relative abundance across depths for each species detected in each mock community group. ZipStrain showed significantly lower depth-dependent variability than MetaPhlAn4 (**paired Wilcoxon signed-rank test, n = 187 species-group pairs, W = 0, p = 1.94 × 10**^**-32**^). **E**, Recovery of expected species from the mock-community reference. ZipStrain recovered a significantly larger fraction of expected species than MetaPhlAn4 (**paired Wilcoxon signed-rank test, n = 93 sample-depth pairs, p = 3.50 × 10**^**-17**^). **F**, Unsupported species calls, defined as detected species not supported by the mock-community reference. ZipStrain produced significantly fewer unsupported calls than MetaPhlAn (**paired Wilcoxon signed-rank test, n = 93 sample-depth pairs, p = 2.90 × 10**^**-17**^). Asterisks denote significance levels; ****, p < 1 × 10^−4^.

For the comparison workflow, we evaluated three approaches: InStrain, standard ZipStrain, and ZipStrain’s “matrix” method, a highly optimized workflow for single-genome comparisons that leverages matrix operations and can run on either CPUs or GPUs (see Methods for details). In single-genome mode, ZipStrain was consistently faster across all approaches: 3× for standard ZipStrain, and 175× and 664× faster for the matrix method on CPU and GPU, respectively, while also using significantly less memory (**Figure 2B**). In all-genomes mode, ZipStrain was 10× faster but consumed more memory; however, ZipStrain dynamically adapts to available memory, reducing RAM usage when necessary (**Figure 2C**). It is also worth noting that ZipStrain computes additional metrics beyond those offered by InStrain, including pairwise gene similarities and the longest identical contiguous stretch.

These performance gains unlock scales of analysis that were previously out of reach. Prior studies using InStrain have typically capped pairwise comparisons at 120 samples, equivalent to 7,140 individual comparisons, with runtimes of over 24 hours, a number that grows quadratically with sample size. With ZipStrain, the same analysis takes just 3-4 hours using the standard mode, under one minute on a MacBook Air M1 in single genome mode, or under 10 seconds on a modern GPU **(Figure 2B)**. Pushing this further, we successfully compared 25,000 samples (over 300 million pairwise computations) for a single genome in approximately 16 hours using one NVIDIA A100 GPU. While this level of comparison would be practically impossible using inStrain (estimated runtime of 50 years), the ZipStrain pipeline is designed for analyses of this magnitude.

#### Consistency of genome presence-absence/relative abundance calculation with ZipStrain

A key aspect of strain-resolved metagenomics is determining which microbes are present or absent in a sample. Although seemingly straightforward, this is complicated by the population structure of microbial genomes (28,29), where strains of the same species can differ substantially in DNA content and unrelated genomes can share stretches of identical DNA. The problem is further exacerbated at low sequencing coverage, where limited data reduces confidence in genome detection. To address this, ZipStrain computes a comprehensive set of genome-level statistics for each genome, including coverage (average sequencing depth across the genome), breadth (fraction of the genome covered by reads), the mean and standard deviation of inter-read gap lengths (distances between adjacent mapped reads), the Breadth-to-Expected Breadth Ratio (BER), and the Fraction of Unexpected Gaps (FUG). The latter two metrics are especially useful for distinguishing true genome presence from spurious read mapping (30): BER captures whether reads are distributed across the genome as expected under random coverage, while FUG detects anomalous clustering of gaps that is indicative of non-specific mapping at low coverages. Together, these metrics enable inference of genome presence or absence using established thresholds (e.g., Metapresence; (30)) or user-defined approaches.

We utilized the experimental design and raw sequencing data from (31) to benchmark ZipStrain’s performance against MetaPhlAn4 (2), a commonly used tool for calculating the relative abundance of microbial taxa. Treichel et al. constructed 13 complex mock communities from up to 70 cultured bacterial strains at set concentrations and sequenced them at depths ranging from 0.1-50 Gbp. Because the ground-truth composition and relative abundances are precisely defined, these mock communities provide an ideal benchmark for evaluating tool accuracy. Additionally, because the samples were physically sequenced rather than simulated *in silico*, the benchmark also captures real-world biases introduced during wet-lab processing, including DNA extraction, library preparation, and Illumina sequencing. ZipStrain achieved significantly higher true-positive rates for detecting expected microbes while substantially reducing false-positive detections of microbes absent from the mock communities (Figure 2E-F). Further, we observed significantly better consistency for relative abundance calculation across different sequencing depths **(Figure 2D)**. These results demonstrate that ZipStrain improves both detection accuracy and abundance estimation robustness relative to MetaPhlAn4.

### Species- and Strain-Level Variation Across a Global Microbiome Dataset

Having established that ZipStrain achieves superior speed, memory efficiency, and detection accuracy relative to existing tools, we next asked what biological insights become accessible at this scale. For the rest of this article, we focus on 2,754 human microbiome metagenomics samples from (32) mapped to two reference databases independently: the DeltaI prokaryotic genome database (14) and the UHGV viral bacteriophage database (33).

#### Genome presence and absence patterns

Genome presence or absence in a sample is typically determined using coverage, breadth, or a combination of both metrics (15,30). ZipStrain provides multiple coverage statistics for each profiled sample, including coverage, breadth, and the breadth-to-expected-breadth ratio (BER), the latter of which is explored in detail in (30). We calculated genome coverage statistics across all samples and genomes in both databases, using the relationship between breadth and expected breadth as the primary metric for presence/absence determination. Notably, bacterial genomes above the Metapresence-recommended BER threshold of ∼0.77 displayed a clearly separable density pattern (**Figure 3A**), highlighting the generalizability of this metric, and potentially this threshold, across different datasets. In contrast, bacteriophage genomes did not show a separation in density and this threshold **(Figure 3B)**. This implies that presence/absence inference in viral datasets may require alternative thresholds or using reference databases more specific to a given environment. Overall, both databases show good representation of genomes for this dataset after filtering for BER <0.77 (genomes per sample for DeltaI and UHGV respectively) **(Figure 3C, D)**.

**Figure 3.**
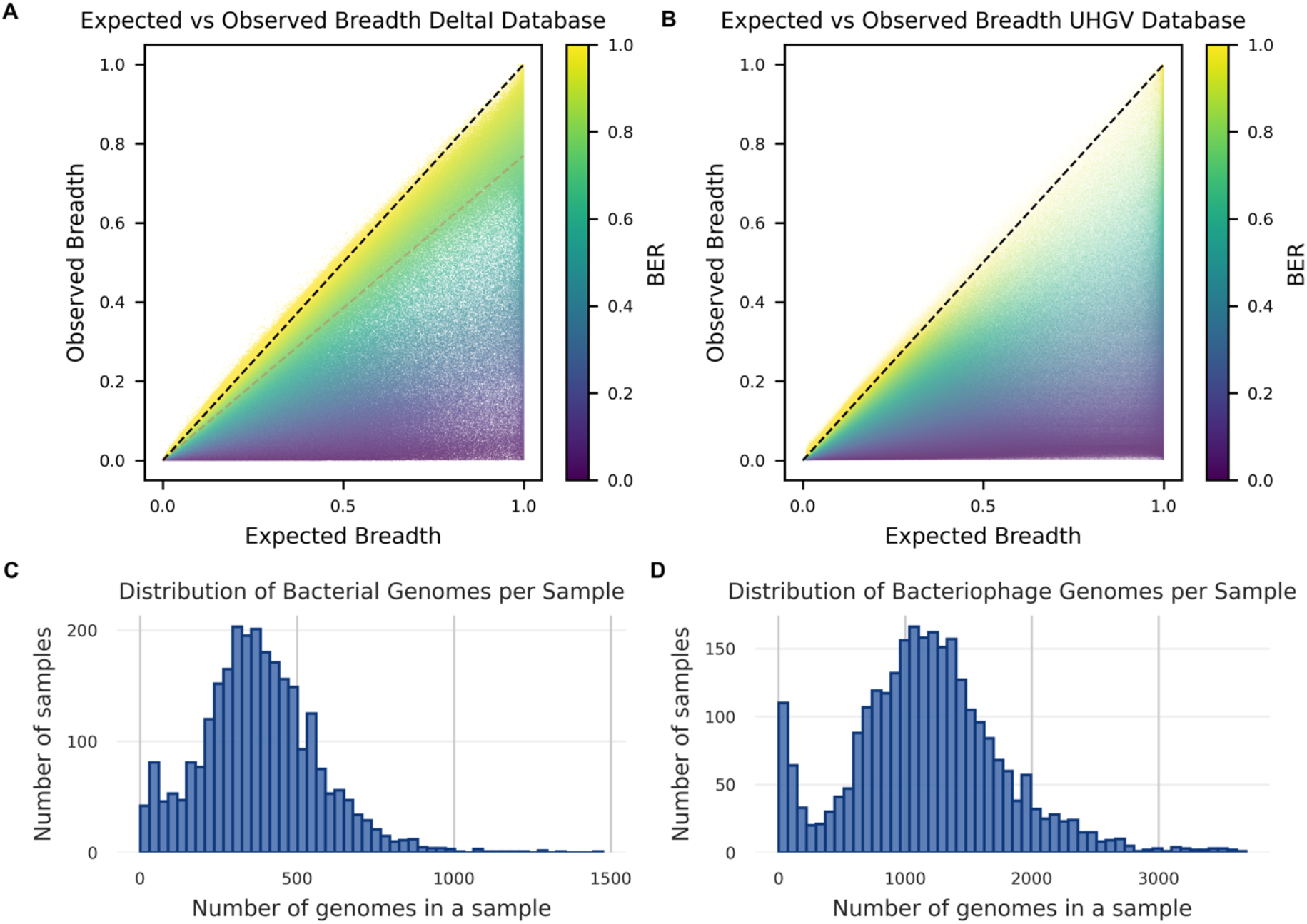
Summary of genome coverage statistics for *(32)*. **(A, B)** Observed breadth vs. expected breadth for all genomes and all samples in this dataset using DeltaI and UHGV as the reference database respectively. Black dashed line shows where observed and expected breadths are the same. The red dashed line in **(A)** shows the threshold suggested by *(30)*. **(C, D)** Histogram showing the distribution of number of genomes detected in samples in this dataset using DeltaI and UHGV as the reference database respectively.

#### Strain Sharing Patterns in Bacteria and Bacteriophages

“Strain sharing rate” is a metric to quantify the extent to which two samples harbor the same microbial strain, and several definitions exist **(Figure 4A)**. The simplest, Strains Shared (SS), counts genomes exceeding a strain-level similarity threshold. Strains Shared over Intersection (SSI), the most commonly used definition (5,32,34), normalizes by genomes detected in both samples. However, this is not biologically meaningful for transmission. Consider a mother harboring 100 species and an infant harboring 20, with 5 detected in both and 4 representing identical strains: SSI returns 80% (4/5), obscuring that only 20% of the infant’s strains and 4% of the mothers are shared. Strains Shared by Each (SSE) captures this asymmetry directly by reporting, for each sample, the fraction of its own genomes shared with the other. We therefore use SSE as our primary strain-sharing metric in all subsequent analyses.

**Figure 4.**
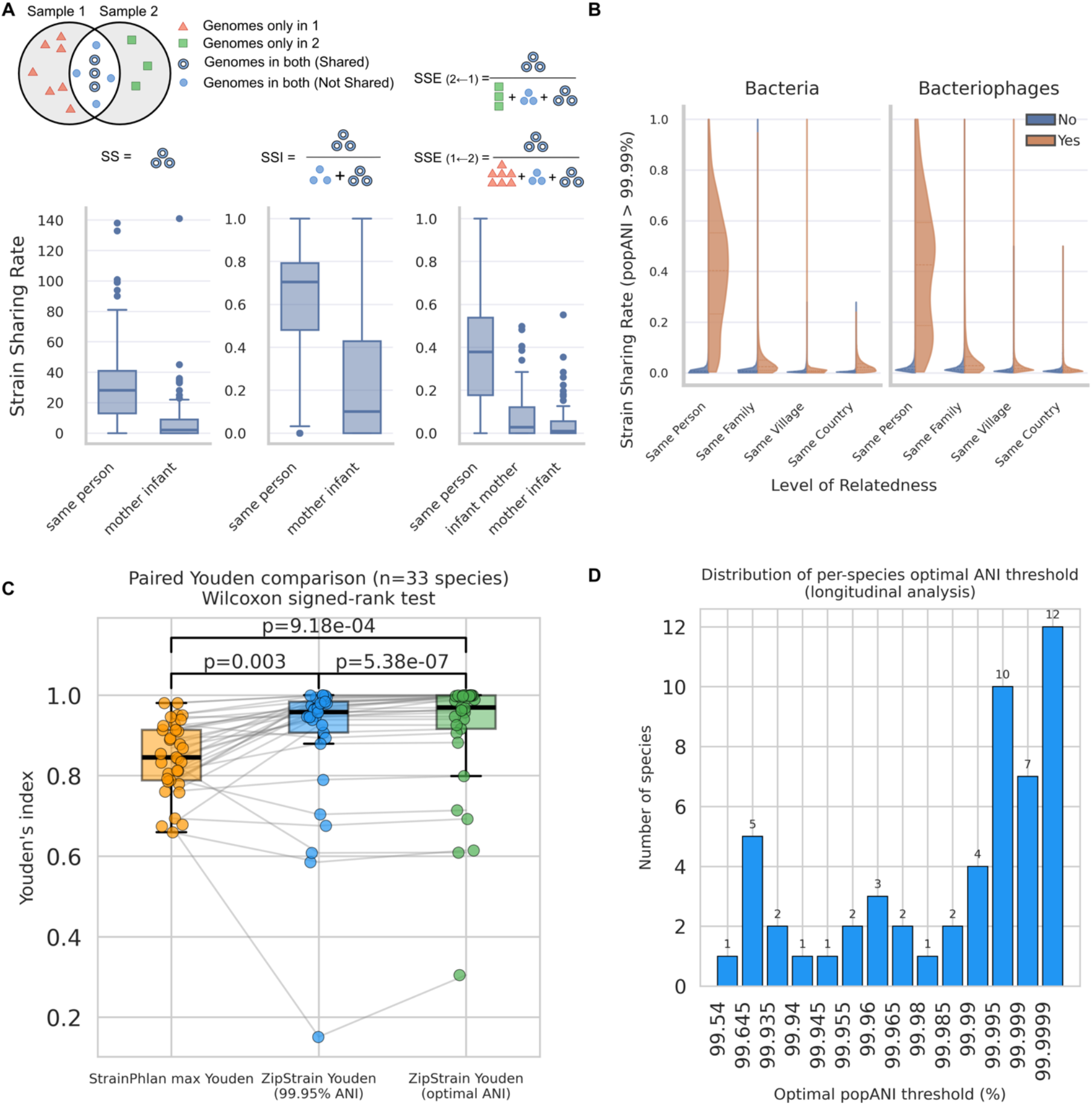
Strain sharing rate calculation with ZipStrain. **(A)** A schematic of three different definitions for strain sharing rate **(B)** Decrease of stain sharing rate (SSE) with the increase of social radius. **(C, D)** Comparing the performance of ZipStrain and StrainPhlAn using time-series within subject samples as ground truth.

Using SSE, we calculated strain sharing rates across four levels of social proximity: (1) the same individual sampled at different time points, (2) different individuals within the same family, (3) different individuals from different families within the same village, and (4) individuals from the same country but different villages **(Figure 4B)**. We observe a clear and significant decline in strain sharing rate as social distance increases, though the difference between the same-village and same-country categories becomes negligible. This trend holds for bacteriophages as well as bacteria. In total, conducting this analysis required performing 4,035,615,045 individual genome comparisons, a scale of analysis that would have been nearly impossible to perform using available genome-wide methods.

We next compared the accuracy of ZipStrain to StrainPhlAn (35), a widely used marker gene– based method that was applied to this dataset (Beghini *et al*. 2021, 2025, Valles-Colomer *et al*. 2023, Ricci *et al*. 2026). In that study, Valles-Colomer *et al*. benchmarked strain-sharing accuracy by comparing the ANI of microbes detected longitudinally within the same individual (assumed to represent the same strain) to the ANI of microbes detected across unrelated individuals (assumed to represent different strains). Performance was summarized on a per-genome basis using the Youden index (defined as sensitivity + specificity − 1). We matched species labels between datasets and compared species-specific Youden index values, finding that ZipStrain significantly outperformed StrainPhlAn across matched species (n=33; *p* = 0.003; **Figure 4C**). This improvement is notable because StrainPhlAn employed species-specific ANI thresholds optimized to maximize the Youden index, whereas ZipStrain used a single global threshold; optimizing ZipStrain’s thresholds in this way improved performance even further (p < 0.001; **Figure 4D**). These results are consistent with prior observations using inStrain (15) and confirm that a genome-wide, population diversity-aware approach provides superior accuracy relative to marker gene-based methods.

#### A Worldwide View of Genome-Wide Similarity Patterns

Perhaps the most compelling functionality of ZipStrain is its ability to perform rapid genome-wide comparisons across large numbers of sample pairs. Here we used our database of over 4 billion pairwise comparisons to introduce a “clonal entropy” metric that captures the degree to which a genome’s pairwise similarity matrix is dominated by one or a few clonal clusters (see Methods for details). Organisms with low clonal entropy form a single dominant cluster, indicative of a highly clonal population **(Figure 5A, D)**, whereas organisms with high clonal entropy are characterized by many small, loosely connected clusters and greater within-species variability **(Figure 5B, E)**.

**Figure 5.**
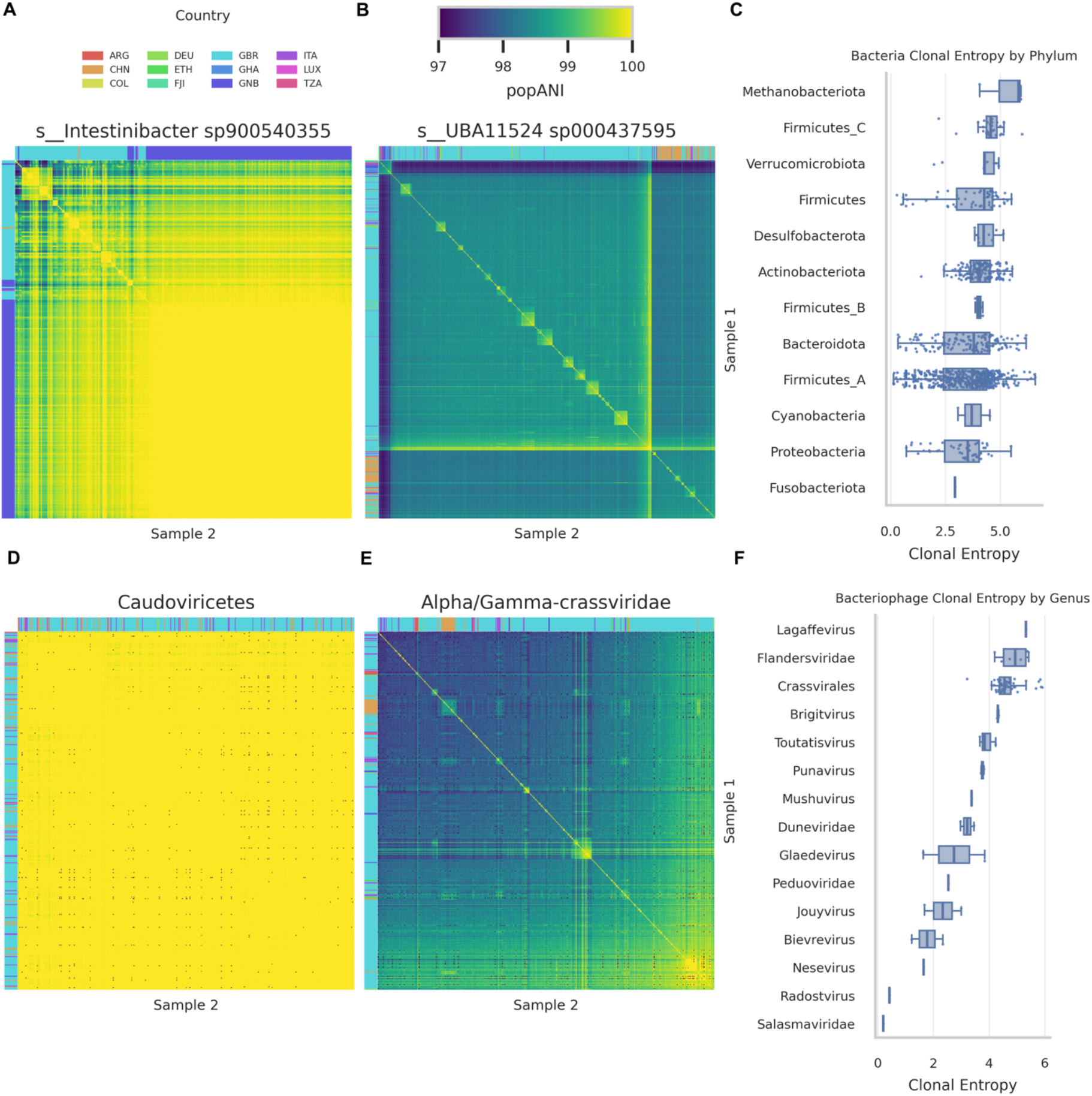
Genome clustering based popANI across all samples. (**A, B**) Clustermaps for genomes with lowest and highest clonal entropy across bacterial genomes respectively. (**C**) Shows the distribution of clonal entropy across different bacterial phyla. (**D, E**) Clustermaps for genomes with lowest and highest clonal entropy across bacteriophage genomes respectively. (**F**) The distribution of clonal entropy across different viral genera.

Applying this metric revealed substantial variation in clonal diversity across the bacteria and bacteriophages in our dataset **(Figure 5C, F)**, with bacteriophages showing consistently higher clonal entropy than bacteria across nearly all genera. Whether clonal entropy differences ultimately reflect transmission route, replication fidelity, or the strength of purifying selection remains to be determined.

#### Gene Divergence Variation in B. infantis

While the genome-level comparison workflow performs comparisons across entire genomes, ZipStrain can also compute ANI between individual genes across all possible sample pairs. This opens the door to a wide range of downstream analyses, including identifying genes whose ANI deviates consistently from the genome-wide average **(Figure 6A)**. Here we demonstrate this functionality by focusing on samples containing *Bifidobacterium infantis*, a keystone colonist of the infant gut microbiome (36–38) as an example. For each gene, we constructed a distribution of deviations from the genome-wide popANI and annotated each gene by its functional category. We identified a diverse range of behaviors across functional categories **(Figure 6A)**. Among genes assigned to COG category J (covering translation, ribosomal structure, and biogenesis), genes with functions directly related to the ribosome or ribosomal RNA displayed positive deviations, while genes involved in tRNA function displayed negative deviations. This type of gene-level analysis provides a finer-grained view of within-species diversity and can highlight functionally meaningful patterns of conservation and variability that would otherwise be obscured in genome-wide comparisons and is possible simply from metagenomics raw reads.

**Figure 6.**
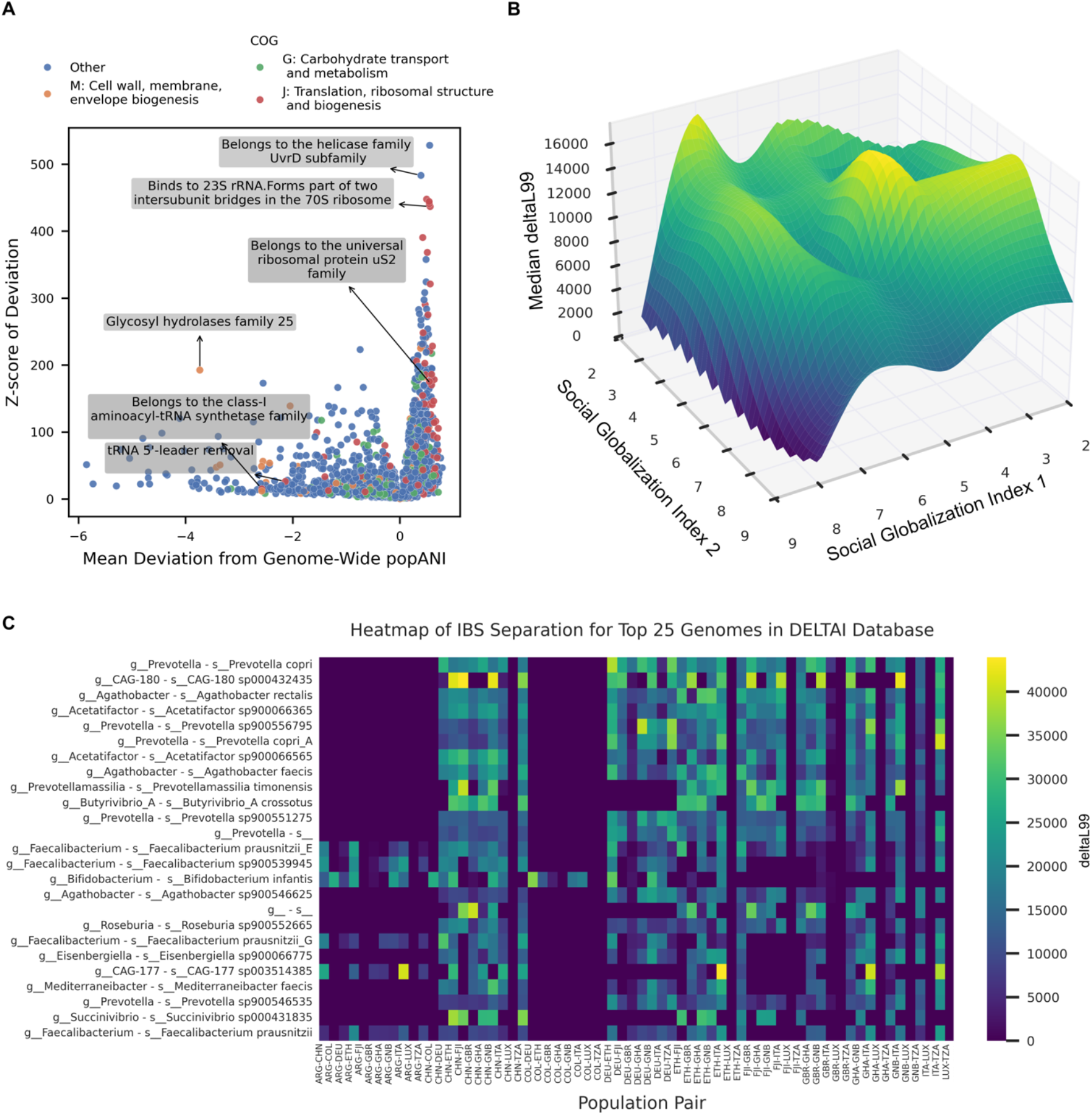
Gene divergence and IBS analysis with ZipStrain. (**A**) Divergence of gene ANI form its genome-wide ANI between sample pairs. (**B**) Relationship between KOFs social globalization score of the sample pair (country of origin) and the median deltaL99. (**C**) Top 25 Median deltaL99 for samples from different country pairs across different genomes, sorted by the median across all country pairs. Three letter codes for countries match those in *(32)*: ARG: Argentina, CHN: China, COL: Colombia, DEU: Germany, ETH: Ethiopia, FJI: Fiji, GHA: Ghana, GNB: Guinea-Bissau, ITA: Italy, LUX: Luxembourg, TZA: Tanzania, GBR: United Kingdom

#### Evolutionary Analysis of Gut Microbial Genomes Across Human Populations

Identity-By-State (IBS) analysis is a metric to measure genetic isolation between bacterial genomes based on the length distribution of identical synonymous sequence tracts shared between genome pairs, where longer tracts reflect more recent gene flow and shorter between-population tracts indicate reproductive isolation (39). We developed a mapping-based implementation of this approach within ZipStrain (21), and here we utilized this approach to study population-level genetic structure of gut microbiome species across human populations **(Figure 6B)**. To quantify connectivity between two countries we used the KOF Social Globalization Index, which assigns a score to each country’s level of social integration with the rest of the world (40). For each country pair, we computed deltaL99 (a measure of evolutionary divergence defined as the difference between within- and between-population L99) for each species detected in both countries. As expected, lower social globalization scores were associated with higher median deltaL99 values, indicating greater genetic isolation between microbial populations from less globally connected countries **(Figure 6B)**. We also examined variation in deltaL99 across microbial species to identify those with the strongest population-level differentiation, a potential signal of long-term co-evolution with human hosts (21). Notably *Prevotella copri* had the highest deltaL99 levels, consistent with its strong association with non-industrialized lifestyles (41), along with several beneficial bacteria including *Bifidobacterium infantis* (**Figure 6C)**. Further work is needed to determine whether these microbes’ associations with human health stem from long-term co-evolution with their human hosts.

## Discussion

The volume of publicly available shotgun metagenomic data has grown exponentially over the past decade (42). Strain-resolved analysis of these data holds substantial promise for understanding patterns of microbial transmission, population structure, and co-evolution with human hosts, but only if the computational tools can keep pace with the data. K-mer-based approaches offer substantial computational efficiency but sacrifice resolution in ways that limit their accuracy in complex or poorly characterized environments. Genome-wide methods are accurate but have historically been too computationally demanding for large-scale analyses. ZipStrain resolves this tension by retaining the biological accuracy of genome-wide comparisons while achieving orders of magnitude speed increases over existing methods.

A central design principle of ZipStrain is flexibility. Sensible defaults and extensive documentation make the tool accessible to users without deep bioinformatics expertise, while the exposure of rich intermediate data and configurable thresholds enable users to adapt the workflow to their specific biological context. The reference genome database can be user-provided, enabling seamless integration with well-curated domain-specific databases such as the UHGG for the human gut (43) or analogous collections for fermented foods (44) and oral microbiomes (45,46). Alternatively, ZipStrain can automatically construct a reference database, extending its applicability to less-characterized environments. The genome presence/absence framework similarly benefits from this design: the BER and FUG metrics performed well for bacterial genomes in the human gut, but presence/absence inference in viral datasets likely requires alternative criteria **(Figure 3)**. By reporting the full distribution of these metrics rather than applying a fixed binary classification, ZipStrain provides users with the information needed to develop context-appropriate thresholds. A similar logic applies to ANI-based strain definitions: while the popANI ≥ 99.999% threshold has been rigorously validated for detecting recent transmission events in human gut bacteria (15), this threshold can be relaxed to study more distant relationships or applied in environmental contexts such as soil or ocean microbiomes where population structure differs substantially.

By increasing the speed of strain-resolved metagenomic analysis by orders of magnitude, ZipStrain goes beyond accelerating existing workflows and enables new analyses that were previously intractable. The clonal entropy metric introduced here **(Figure 5)** is a direct product of this new capability. Characterizing clonal structure meaningfully requires pairwise comparisons across thousands of samples, and the striking variation in clonal diversity we observe across bacterial and bacteriophage taxa would have been inaccessible with prior genome-wide methods. ZipStrain also introduces new features whose utility will extend across diverse research applications. Gene-level divergence analysis (identifying individual genes whose ANI consistently deviates from genome-wide expectations; (**Figure 6**)) offers an ability to detect ongoing selection, horizontal gene transfer, or functional diversification directly from metagenomic reads. The IBS framework provides a rigorous, mapping-based method for quantifying evolutionary isolation between microbial populations, with direct relevance to questions of host-microbe co-evolution and the biogeography of human-associated microbes.

Several limitations of ZipStrain are worth noting. Like all reference-based metagenomic approaches, ZipStrain is constrained by the completeness of the reference genome database; taxa absent from the database will not be profiled. ZipStrain does not currently account for genomic insertions and deletions when computing ANI, which may underestimate divergence between strains that differ substantially in gene content or harbor large mobile genetic elements. Finally, in the interest of performance, ZipStrain does not implement nucleotide divergence, linkage disequilibrium analysis, and other features available in inStrain that can be useful for profiling within-population recombination and selection.

## Conclusions

ZipStrain enables accurate, genome-wide strain-resolved metagenomics at a scale that was not previously achievable with existing tools. Benchmarking demonstrates consistent order-of-magnitude speedups in both profiling and comparison, with the largest gains realized when comparing large numbers of samples against comprehensive reference databases. Accuracy assessments confirm that the genome-wide, population diversity-aware approach of ZipStrain yields superior strain discrimination relative to marker gene-based methods. Applied to a global cohort of 2,754 human gut metagenomes, ZipStrain reveals that strain sharing tracks social proximity across both bacteria and bacteriophages, that clonal structure varies substantially across taxa in ways that may reflect fundamental differences in microbial lifestyle and transmission, and that population-level evolutionary isolation of gut microbes covaries with human social connectivity. ZipStrain is distributed as an open-source Python package with a companion Nextflow pipeline, comprehensive online documentation, and tutorials, facilitating reproducible deployment across diverse computing environments. By removing the computational barriers that have constrained strain-resolved analysis to small sample sizes, ZipStrain enables users to leverage the full scale of available metagenomic data to address questions about microbial transmission, evolution, and ecology at global resolution.

## Declarations

### Ethics approval and consent to participate

This study used only publicly available, de-identified metagenomic sequence data. No new human or animal data were collected.

### Consent for publication

Not applicable.

### Availability of data and materials

The metagenomic sequencing data analyzed in this study are publicly available from the European Nucleotide Archive under accession numbers provided in (32). The DeltaI prokaryotic genome database is available as described in (14). The UHGV bacteriophage database is available as described in (33). ZipStrain is available as an open-source Python package at https://github.com/OlmLab/ZipStrain under an MIT license.

### Authors’ contributions

PG developed ZipStrain, implemented all software components, and performed all computational analyses. MRO conceived and supervised the project and wrote the manuscript. JBE provided intellectual guidance and critical feedback throughout the project. All authors read and approved the final manuscript.

## Acknowledgements

The authors thank members of the Olm Lab for testing the software during development.

This work utilized the Blanca condo computing resource at the University of Colorado Boulder. Blanca is jointly funded by computing users and the University of Colorado Boulder.

## Methods

### Metadata preparation

For analyses presented in this manuscript, we entirely utilized published datasets with (32). Specifically, we utilized a subset of 2754 samples that were assigned family IDs and had highest sequencing depths (**Supplementary Table 1**).

### Database preparation

For bacterial genomes, we used DeltaI genome database (14). The genes for this database were extracted using prodigal v2.6.3. For the bacteriophage database, we used the UHGV High-quality representative genomes. We extracted the viral genes for these vOTUs using prodigal-gv v2.11.0 (47,48)

### Mapping reads to the reference database

For mapping reads to both reference database we used bowtie2 with default settings and discarded any unmapped reads using “samtools view -bS -F 4” samtools command.

### Profiling

In order to generate ZipStrain profiles, we utilized “from_sra_to_profile” workflow in ZipStrain’s Nextflow pipeline that when provided the SRA accession for a sample, it downloads the sample from SRA using SRAToolkit, maps the reads to the reference genome using bowtie2, filters unmapped reads, and finally runs ZipStrain’s profile-single command on the “.bam” files.

This step generated two files for each sample: profile and genome_statistics tables both in “.parquet” format.

### Comparison

#### Comparison with ZipStrain’s compare workflow

To create the genome comparison tables, we performed pairwise comparisons at genome-level using “compare_genomes” workflow with min_cov = 5, min_gene_compare_len=200 and engine=“polars”. Comparison workflow utilizes Polars (19) or DuckDB (18) Python libraries to perform inner join between the profile pairs on scaffold, position, and genome columns. After joining, SNPs are identified by comparing the nucleotide frequencies at matching loci, see https://olmlab.github.io/ZipStrain/api/#compare. As an example, for popANI if the dot product of [A, T, C, G] frequencies is zero between the two profiles, then a popSNP is identified at that location. Genome-wide and gene-wide ANIs are calculated by dividing the number of non-SNP locations by the total overlapping region for every genome and gene respectively. Additionally, genome comparison workflow, keeps track of the longest identical stretch for every genome between two samples by identifying the longest stretch on the genome that is covered in both samples and is not interrupted by a SNP.

#### Comparison with ZipStrain’s matrix workflow

First, ZipStrain’s “.parquet” profiles are indexed to form a contiguous matrix for each genome. The result of this step is a “HDF5” file that holds the matrices for all samples for the genome of interest. This is done using “zipstrain utilities build-matrix-db” command. Once this step is over, the comparisons can be executed using “zipstrain utilities matrix-compare” command. In the matrix method, ANI and IBS calculations are defined as matrix operations and PyTorch (20) is used to perform these operations in an efficient manner on a CPU or GPU depending on the availability.

### Strain sharing

Given the lack of standard definition for strain sharing rate, we defined three different definitions and calculated each for mother infant pairs in our dataset: Strains Shared (SS): This metric simply counts the number of the strain sharing events. A strain sharing event between two samples is when the covered region in both samples in a pair (here 5x coverage) exceeds a threshold (here 50 kbp for bacterial genomes and 10 kbp for bacteriophage genomes) and the calculated genome-wide popANI is above a defined threshold (here 99.99).

Strains Shared over Intersecting genomes (SSI): This metric is defined as the number of strain sharing events between two samples over the number of genomes that exists in both samples. For this metric we need to define a way to call a genome present in a sample for strain sharing calculations. We call a genome present in a sample if that genome is covered at least more than the overlap threshold (more than 50 kbp at 5x for bacterial genome and more than 10 kbp at 5x coverage for bacteriophage genomes). Once the genomes present in each sample is determined, say *G*_*A*_and *G*_*B*_ represent set of genomes in sample A and B respectively, then SSI can be calculated by:

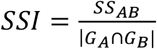

Strains Shared by Each (SSE): This definition of strain sharing assigns a rate to each sample in a pair. Using the previous notations, SSE can be computed for each sample in a pair by:

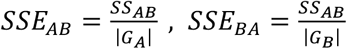

We find this definition of strain sharing rate particularly appealing because of the directional nature of the metric.

### Clonal entropy

For all the ANI analysis in this study, we utilized the comparison tables. Each row in the comparison table reports different metrics. For analysis based on genome-wide ANI we used sample_1, sample_2, genome_pop_ani. Prior to any clustering, we filtered the comparisons to only include genome comparisons where the coordinates covered in both samples exceeds 50 kilo base pairs (kbp) for bacteria and 10 kbp for bacteriophages.

For clonal entropy calculations, we excluded sample pairs that are originating from the same individual. Next, we assigned clusters to each sample by setting 99.9% similarity as the clustering threshold and using “average” linkage mode in SciPy (49)to perform clustering and using distance as the clustering criterion. As a result, for every genome, each sample pair with popANI>99.9% the samples will be assigned the same clonal cluster.

To calculate clonal entropy, we first ruled out any rare genome that did not exist in more than 100 samples. Next, for each genome we calculated the Shannon index by using the relative abundance of the clonal clusters as an estimator of the probability of observing the cluster, similar to alpha diversity calculation, and calculated the clonal entropy by:

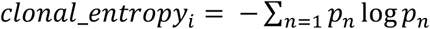

Where:

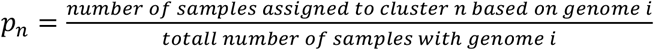

Finally, we sorted the genomes by their clonal entropy metric and generated cluster maps in seaborn with SciPy, using the same filtering criteria mentioned above, for genomes with lowest and highest clonal entropy in both bacterial and bacteriophage reference database reported in **Figure 5**.

### Identity-By-State IBS

This analysis is a mapping-based alternative to the analysis introduced by (21). The genome comparison table reports the longest identical stretch for every genome in a sample pair in the column “max_consecutive_length”. For IBS calculations, we first assign each sample to a population (here countries). Then we consider the desired population pairs and build two cumulative probability distributions for each population pairs based on the “max_consecutive_length” values; One for pairs where both samples belong to the same population and one for pairs where each sample is coming from one of the populations in the popluation pair. After building these two cumulative distributions, we find the length for which the CDF>99% in both within and between population distributions (within_populationL99 and between_populationL99). We denoted the difference between these two quantities as deltaL99. This operation yields a value for each genome for each population pair. DeltaL99 was used to generate Figure 6 B, C. For Figure 6 B, we utilized the mean KOF’s social globalization indices and averaged the values between years 2015-2020 (Assuming this is when the sample collection is happened) for every country.

### Benchmarking ZipStrain

For the performance benchmark, we first identified the genome with the highest sample presence using a minimum coverage threshold of >1x and a breadth-of-coverage threshold of 0.8, which selected “*GUT_GENOME251083*.*fna*” for the single-genome comparison. For the all-genome benchmark, samples were ranked by richness to ensure that selected cohorts contained sufficient genomes for all-genome comparisons. We then selected the top 100 samples and generated subsets of 2, 5, 10, 50, and 100 samples, with three random replicates for each subset size except for the full 100-sample set. These subsets were profiled and compared using ZipStrain v0.9.10 and InStrain v1.10.0 on a CPU node with 10 CPUs and 250 GB RAM. Where GPU is mentioned we utilized NVIDIA A40 available on the Alpine supercomputing cluster.

For the taxonomy benchmark, mock-community sequencing samples from (31) were downloaded from the Sequence Read Archive (SRA) and profiled with ZipStrain agains custom database made by sylph for each sample independently and MetaPhlAn4 using MetaPhlAn database version “mpa_vJun23_CHOCOPhlAnSGB_202403”. MetaPhlAn per-sample abundance tables were merged across samples, and MetaPhlAn taxonomy identifiers were converted to GTDB-compatible notation using “sgb_to_gtdb_profile.py” provided in MetaPhlAn4’s GitHub repository. The resulting profiles were compared against the reference metadata (31). Robustness to sequencing depth was quantified for each species detected in a sample across at least three depths as the standard error of relative abundance across depths, defined as standard deviation / sqrt(n), where n is the number of depths at which the species was detected. Species recovery was calculated as the fraction of expected reference species detected in each sample, and unsupported calls were defined as detected species not supported by the reference metadata after permissive GTDB-compatible taxonomic matching.

### LLM disclosure

Portions of this manuscript were edited with the assistance of Claude (Anthropic), a large language model (LLM). All scientific content, analyses, and conclusions were generated and verified by the authors.

